# COCONUT: An analysis of coiled-coil regions in proteins

**DOI:** 10.1101/2024.03.25.586698

**Authors:** Neelesh Soni, M. S. Madhusudhan

## Abstract

**Motivation:** The molecular rules determine the strength and orientation (parallel or antiparallel) of interacting coiled-coil helices in protein-protein interactions. Interpreting these rules is crucial for identifying novel protein-protein interactions, designing competitive binders, and constructing large assemblies containing coiled-coil domains. This study establishes the molecular principles that dictate the strength and orientation of coiled-coil interactions, providing insights relevant to these applications.

**Results:** We examined how hydrophobic contacts determine structural specificity within coiled-coil dimers. Our analysis revealed that the hydrophobic core densities differ between parallel and antiparallel dimer confirmations, highlighting their importance in stabilizing different structural arrangements. We developed **CO**iled-**CO**il a**N**alysis **UT**ility (COCONUT), a computational platform with machine learning models, validated for predictive capabilities in various applications. Using COCONUT’s pipeline for coiled-coil analysis and modeling, we predicted the orientation of substitution-sensitive coiled-coil dimer, identified residue pairings in non-canonical coiled-coil heterodimer, and constructed *n-stranded* coiled-coil model. These results demonstrate COCONUT’s utility as a computational framework for interpreting and modeling coiled-coil structures.

**Availability and implementation:** COCONUT is an open-source and free Python package available here https://github.com/neeleshsoni21/COCONUT. The documentation is available in the source code and here: https://neeleshsoni21.github.io/COCONUT/

## 1 Introduction

Coiled-coils (CC) are diverse structural motifs found in 10% of all proteins and serve pivotal roles in molecular recognition, protein folding, and signal transduction (1–5). The primary function of coiled-coils is to provide structural integrity and resilience to protein assemblies by forming stable homo- or hetero-oligomeric complexes (7–11). About 80% of all coiled-coils are hetero-oligomers and play a significant role in quaternary interactions (6). The specificity of protein-protein interactions is achieved through coiled-coils involving two or more alpha-helices twisted around each other in a left-handed supercoil (12–15). The characteristic heptad sequence repeat pattern (a-b-c-d-e-f-g) shapes their overall structure, with the residues at heptad positions ‘a’ and ‘d’ forming a buried interface, while residues at other positions are exposed to the solvent (15, 16). Through this design, a range of molecular forces (e.g. hydrophobic contacts and salt bridges) emerge, guiding the stability of coiled-coils. Coiled-coil stability refers to the strength or propensity of a coiled-coil structure to maintain its helical conformation. This facilitates specific residue pairing, enabling coiled-coils to have a capacity for molecular recognition (14, 17).

Several software programs and databases have been streamlined for characterizing coiled-coil motifs based on protein sequences (18–32). These resources aid in identifying coiled-coil motifs and predicting their location and sizes in protein sequences (33). However, the existing tools do not predict the complementary sequences that could form a coiled-coil structure, especially in hetero-oligomers. These details are crucial for understanding their function, structural modeling, and designing new molecular tools (33–35).

Various studies have attempted to characterize the stability of coiled-coils using techniques such as mutagenesis (40–42, 49, 50), the effect of pH (43–46, 51), and the role of hydrophobic interactions (to name a few) (20, 47, 48). While these studies have undoubtedly advanced our understanding of coiled-coils, we do not yet have the means of using this knowledge in a predictive sense, i.e., identifying potential binding partners of coiled-coils, predicting coiled-coil orientation, etc.

To address this issue, we have developed **CO**iled-**CO**il a**N**alysis **UT**ility (COCONUT), an open-source coiled-coil scoring and modeling platform. We describe the underlying dataset, preprocessing methods, statistical frameworks, and the development of a Random Forest-based predictive model to deduce the molecular determinants of stability. Given coiled-coil dimer forming sequences, COCONUT can deduce the best amino acid pairing and predict its orientation (parallel or antiparallel dimer).

This capability can be used in identifying novel protein interactions, designing competitive binders to coiled-coil proteins, building coiled-coil models of large assemblies, and, generally, any application that requires stable-specific coiled-coil interactions. In addition to our previously published application (53), we demonstrate COCONUT’s predictive capabilities using three diverse modeling scenarios. First, we employed a canonical coiled-coil leucine zipper sequence to confirm COCONUT’s proficiency in precisely determining dimer orientation. This resulted in an alignment closely matching the known crystal structure with minimal root-mean-square deviation (RMSD). Subsequently, we broadened COCONUT’s application to a non-canonical Keratin K5-K14 dimer. Despite lacking training data for this dimer class, COCONUT successfully generated models that aligned well with the known structures. Finally, we modeled a coiled-coil trimer domain of Tumor necrosis factor receptors, showcasing its precision in the structural predictions of *n-stranded* coiled-coils.

## 2 Methods

### 2.1 Coiled-coil Dataset and Preprocessing

The coiled-coil structures analyzed in this study were retrieved from the CC+ database (22, 52). We selected dimers with a sequence identity of less than 35% and a minimum and maximum number of amino acids as 11 and 100. We obtained 1149 coiled-coil dimers containing 252 parallel and 897 anti-parallel orientations.

### 2.2 Extracting coiled-coil dimer segments

Knob-into-hole packing is a characteristic feature of coiled-coil structures, where specific amino acid residues from one helix fit into spaces created by the surrounding residues of an adjacent helix, much like a knob fitting into a hole (**Fig. 1A, B**) (12, 13, 15). To extract all residue pairs that form a Knob-into-hole interaction, a sliding window along the helix is used to identify sets of residues (hex pair) in physical proximity and categorized as “dad”/“ada” hex pairs (**Fig. 1C**). A “dad” hex pair consists of twelve amino acids (six from each helix), positioned at heptad a-a’, flanked by d-d’ and e-e’/g-g’ pairs (**Fig. 1B**). Conversely, the “ada” hex pair comprises twelve amino acids at heptad d-d’, flanked by a-a’ and e-e’/g-g’ pairs. For instance, for a parallel dimer, a “dad” pair consists of heptad positions 1d, 1d’, 2a, 2a’, 2d, 2d’, 1e, 1e’, 1g, 1g’, 2e, and 2e’ (**Fig. 1C**). These residues are numbered according to their position in the heptad repeat, which runs from the beginning to the end of the coiled-coil dimer. A similar hex pair assignment is done for the anti-parallel dimers (**Fig. 1C**). The previously selected coiled-coil dimer dataset (1149 coiled-coil dimers) provides 1516 parallel hex pairs, and 3424 anti-parallel hex pairs (**Fig. 1D, E**). This hex pair dataset was used to deduce structural features for further analysis.

**Figure 1:**
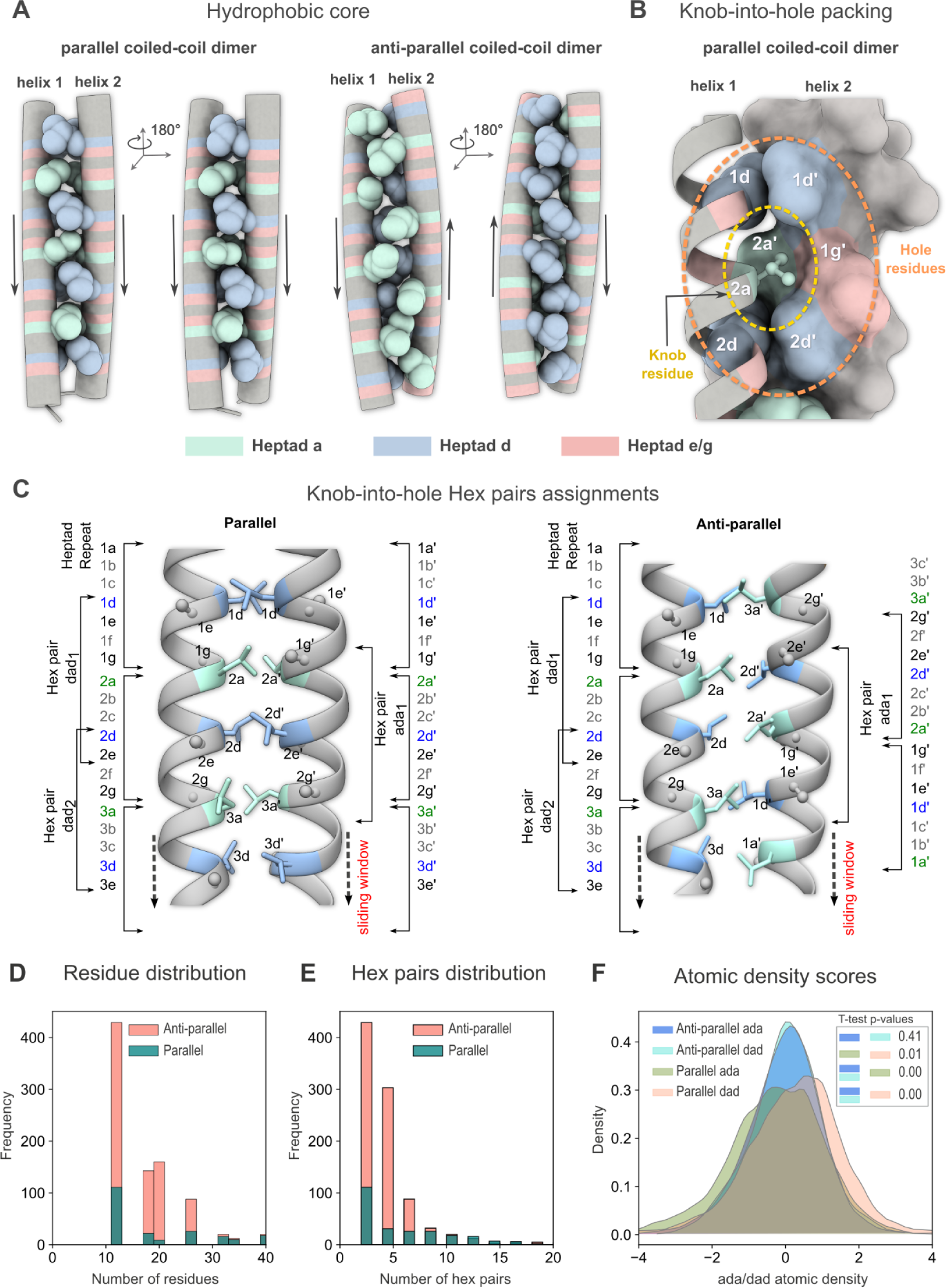
Coiled-coil interactions and knob-into-hole packing. (A) Position of the “hydrophobic core cylinder” in a parallel (PDB 2ZTA) and antiparallel (PDB 1FXK) coiled-coil dimer formed by hydrophobic atoms from residues at heptad positions a/a’ (turquoise) and d/d’ (light blue). Side chains of amino acids at these positions clipped at C_δ_ are shown in a sphere representation. Hydrophilic residues occupy positions at heptad repeat e/e’ and g/g’ (salmon) that form salt bridges across helices. Each helix is shown in a tube representation with vertical arrows highlighting the direction of the amino acid sequence from the N-terminus to the C-terminus. For parallel coiled-coil dimers, hydrophobic atoms at heptad positions ‘a’ and ‘d’ alternate on both sides of the hydrophobic core. Hydrophobic atoms at heptad positions ‘a’ or ‘d’ remain on either side of the hydrophobic core for the antiparallel dimer. (B) A knob-into-hole packing example in a parallel coiled-coil dimer. A hole is a hydrophobic pocket generally formed by a set of six residues (white labels) by their hydrophobic side chain atoms (orange dotted circle). This hole provides an enclosure to the hydrophobic side chain of the knob residue (yellow dotted circle). (C) A parallel and antiparallel coiled-coil dimer with hydrophobic core residues in stick representation. A hex pair is a set of six hydrophobic core residues and six hydrophilic residues that possibly interact in a Knob-into-hole packing. Hex pairs are of two types based on heptad repeats: Hex pair “dad” and “ada”. Hex pairs of a coiled-coil dimer are extracted by linearly sliding through the hex pairs along the coiled-coil axis. (D) Frequency distribution of the number of residues in anti-parallel and parallel dimers. (E) Distribution of hex pairs in anti-parallel and parallel dimers. (F) Kernel density estimation plots of atomic density scores, comparing anti-parallel “ada”/“dad” distributions with parallel “ada”/“dad” distributions.

### 2.3 Features of coiled-coil structures

To deduce the molecular determinants of stability and specificity of coiled-coil structures, we define various parameters that capture the molecular interactions and their differences between the parallel and anti-parallel coiled-coils. Below is the list of features and their definitions that were used subsequently in the classification and stability prediction.

#### 2.3.1 Amino acid pair preferences for heptad positions

Using the hex-pairs dataset, we extracted the pairwise frequency of amino acids at different heptad pairs. We categorized the heptad pair interactions into three types: 1) heptad pairs (a-a, d-d, a-d) that form the hydrophobic core of a coiled-coil, 2) heptad pairs (a-g, d-e, d-g, a-e) that support the hydrophobic core and prevent water from penetrating the core, 3) heptad pairs (e-g, e-e, g-g) that form salt bridges between the two helices in a coiled-coil dimer. We calculated the pairwise amino acid frequencies for each heptad pair and normalized them by their total number. These distributions provide probability estimates of amino acid pairs given the heptad pair position across parallel (**Fig. 2A**) and anti-parallel dimers (**Fig. 2B**).

**Figure 2:**
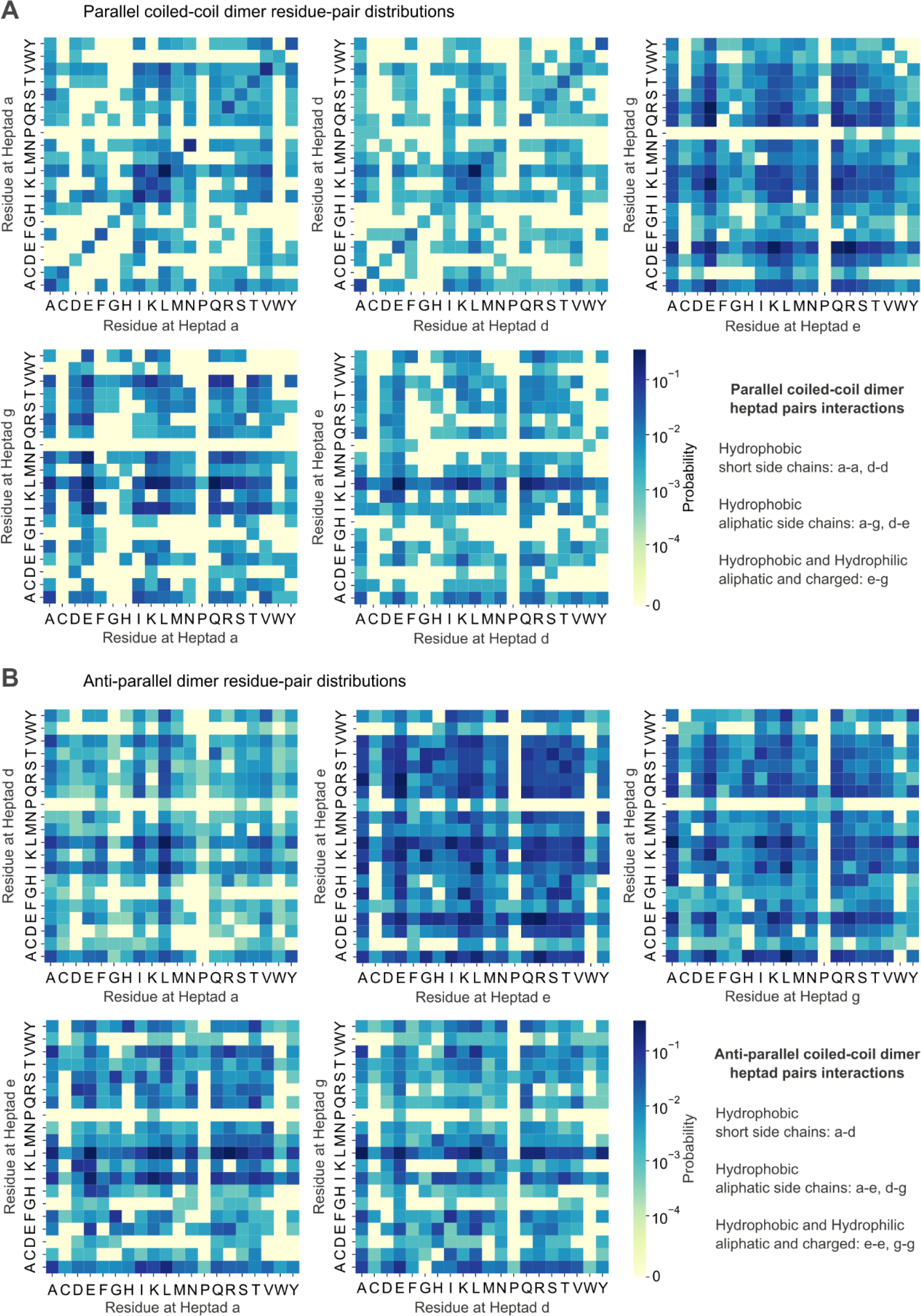
Amino acid frequency distribution of heptad pairs in coiled-coil dimers. Heatmaps of residue-pair distributions for coiled-coil dimers, separated into two categories: parallel dimers (A) and anti-parallel dimers (B). Each heatmap corresponds to different positions within the heptad repeat (‘a’ to ‘g’), with the axes representing the 20 standard amino acids. The color intensity indicates the probability of occurrence between pairs of residues at specified positions, ranging from high (dark blue) to low (light yellow). These distributions reflect the propensity of amino acid pairs to interact based on their position within the heptad repeat and categorized by the nature of side chains: hydrophobic with short side chains, hydrophobic short and long aliphatic side chains, and combinations of hydrophobic, hydrophilic side chains, thereby providing insight into the structural preferences that govern coiled-coil dimer formation.

#### 2.3.2 Atomic density scores for hydrophobic core

We established pairwise scores for residues engaged in physical interactions within the hydrophobic core to quantify the local packing of atoms. These atomic densities were estimated within the hydrophobic core, formed by residues at the heptad pairs a-a’ and d-d’. Each amino acid was assigned a score calculated as the total number of non-hydrogen atoms at β and γ positions in the side chains (**Table 1**). Amino acids with the following atoms were given additional scores as indicated: C^γ^ sp^2^ hybridized carbon = +1, S^γ^ = +1.5, S^δ^ = 0.5, C^δ^ = sp^3^ hybridized carbon = +0.25. These additional scores were informed from the molecular properties and account for the bulkiness/size (such as an S atom being bulkier than atom C) and degrees of freedom (for sp^2^ hybridization, planar molecule). All other atoms at C^δ^ and beyond were ignored, as they do not contribute to the hydrophobic core of the coiled-coil dimer. A pairwise score (PS) of heptad pairs at a-a (*PS*(*a* − *a*)), d-d (*PS*(*d* − *d*)), and a-d (*PS*(*a* − *d*)) was determined by adding the individual scores of amino acids at these positions. The pairwise scores were used to calculate the deviation at a given heptad pair (atomic density scores i.e., *ADS*) compared to the successive and preceding heptad pairs. For instance, a *ADS* score at the heptad d-d’ position *ADS*(*ada*_1)_ can be calculated by taking the mean of the *PS* scores at successive and preceding a-a’ positions and subtracting the *PS* scores of the current position d-d’. The equations below (**Eq. 1 to 4**) illustrate the computations of *ADS* scores for hex pairs “ada” and “dad” (**Fig. 1C**) for both parallel and anti-parallel dimers.

**Table 1:** Atomic density scores for individual amino acids. Every amino acid was assigned a constant score representing its contribution towards the atomic density score at the hydrophobic core of a coiled-coil dimer (**Methods 2.2.2**)

|  |  |  |  |  |  |  |  |  |  |
| --- | --- | --- | --- | --- | --- | --- | --- | --- | --- |
| A | C | D | E | F | G | H | I | K | L |
| 1 | 3.5 | 3 | 2 | 3 | 0 | 3 | 3.25 | 2 | 2.5 |
| M | N | P | Q | R | S | T | V | W | Y |
| 2.53 | 3 | 4 | 2 | 2 | 2 | 3 | 3 | 3 | 3 |

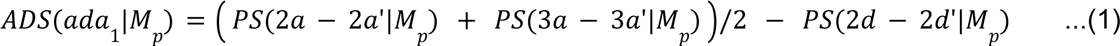

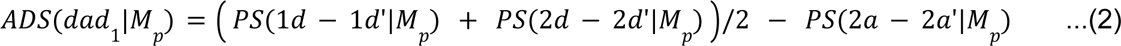

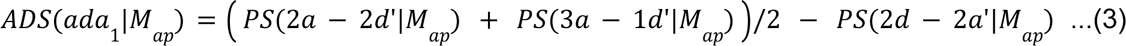

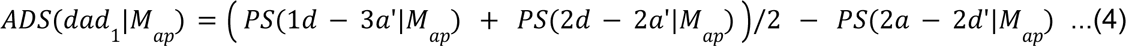

Where *PS*(*a* − *a*’|*M*_*p*_) and *PS*(*d* − *d*’|*M*_*p*_) denote pairwise scores at heptad pair a-a’ and d-d’ for parallel dimers, *PS*(*a* − *d*’|*M*_*ap*_) and *PS*(*d* − *a*’|*M*_*ap*_) denote pairwise scores at heptad pair a-d’ and d-a’ for antiparallel dimers.

To model the ADS distributions as smooth estimates of a probability density function, we utilized the kernel density estimator (55–57) with ‘scott’ method (56) for bandwidth estimation. This approximates the ADS score distributions for “ada” or “dad” hex pairs within parallel and anti-parallel dimer configurations (**Fig. 1F**). Following the training phase, we conducted comparative analyses of the resulting distributions to verify their statistical differences in classifying the hex pair types. Since the distributions were derived using a ‘Gaussian’ kernel, their comparisons were made via a two-tailed T-test at a 95% confidence interval (**Fig. 1F-inset**). These distributions were utilized to generate probability estimates for new data points while making predictions (**Fig. 1F**).

### 2.4 Building the classifier for predicting structural stability

We adopted a machine-learning approach to analyze and predict the structural stability of coiled-coil motifs by utilizing information from frequency distributions and atomic density score distributions. These extracted features were assembled into vectors with appropriate padding to standardize their lengths across different instances. Subsequently, we created a dataset designated for training and validation purposes, which facilitated the development of a predictive model. We trained a Random Forest Classifier (58, 59) with stratified k-fold cross-validation, executing partitions at 2, 5, and 10-fold (**Fig. 3A, B, C**). During the training, 100 estimators were used with bootstrapping and splitting the nodes with entropy criteria until all leaves were pure. The output of the models is the prediction of the dimer’s orientation, which was compared against the known labels. The cross-validation process quantitatively summarized the average prediction accuracy of the dataset along with Matthew’s Correlation Coefficient (MCC) (60) and confusion metrics (**Table 2**). Since MCC does not change significantly at various fold cross-validations, 10-fold cross-validated trained models were used for further predictions and analysis (**Fig. 4**).

**Figure 3:**
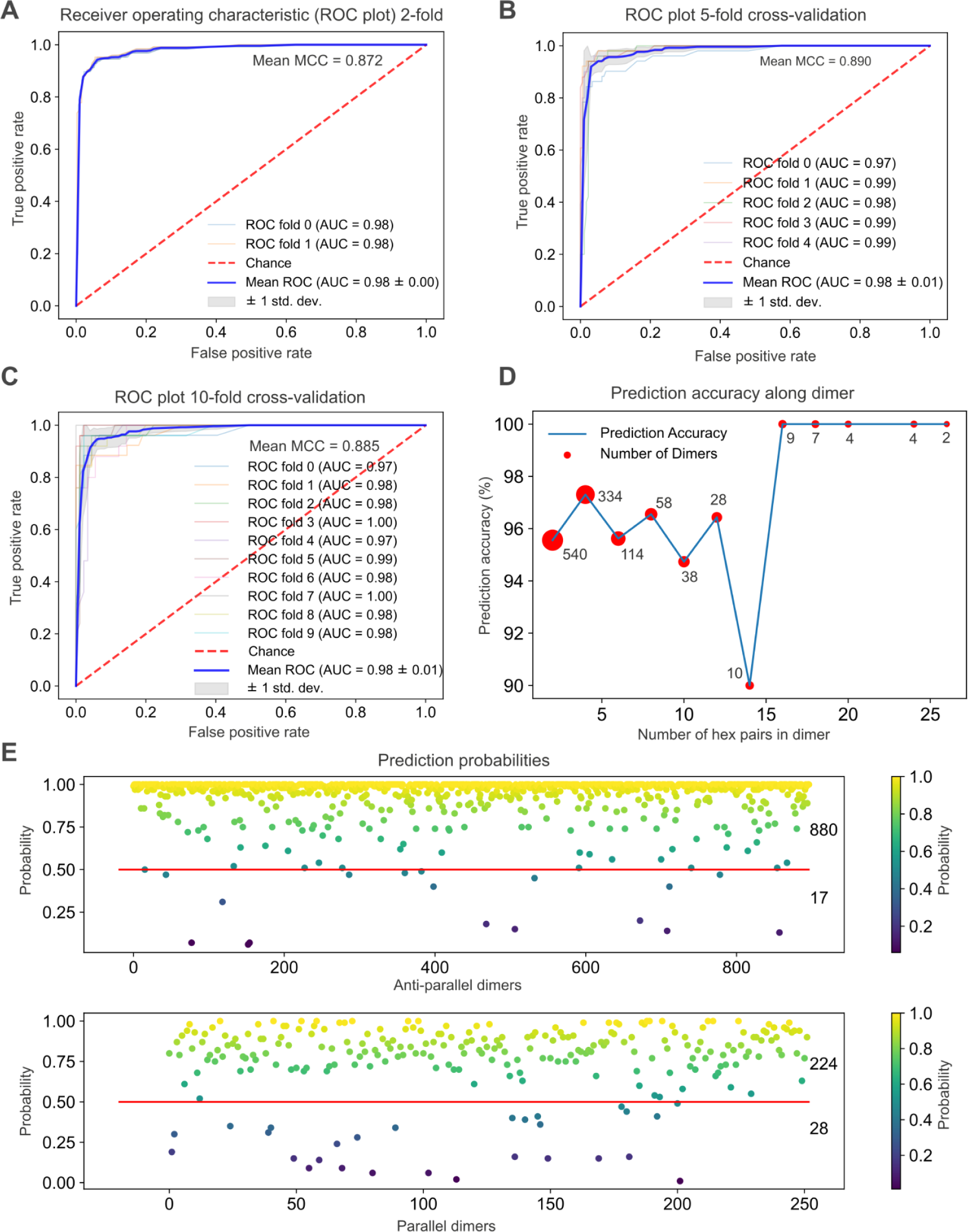
Evaluation of COCONUT’s predictive performance. (A) ROC plot for 2-fold cross-validation displaying true positive rate vs. false positive rate, with a mean Matthews Correlation Coefficient (MCC) of 0.872. (B) ROC plot for 5-fold cross-validation with a mean Area Under the Curve (AUC) of 0.98 and a mean MCC of 0.890. (C) ROC plot for 10-fold cross-validation, indicating a mean MCC of 0.885. Individual ROC curves for each fold are shown, highlighting the consistent AUC values across folds. (D) Graph showing the relationship between prediction accuracy and the number of hex pairs in a dimer, with the size of the bubble representing the number of dimers at each data point. (E) Scatter plots of prediction probabilities for anti-parallel (top) and parallel (bottom) dimers. Each point represents a dimer, color-coded by probability, with the red line denoting the threshold for classification. The number of dimers accurately predicted (above the threshold line) and misclassified (below the threshold line) is noted at the right end of each plot.

**Table 2:** Results from 2/5/10-fold cross-validations of Random Forest Model. The table provides confusion matrices along with Matthew’s Correlation Coefficient (MCC) and accuracy (ACC) for individual sets comprising antiparallel (AP) and parallel (P) dimers from 2/5/10-fold cross-validations (CV) of the Random Forest Estimator. (**Methods 2.4**). The last row provides the mean MCC and average accuracy overall sets in each fold validation.

|  | 10-Fold CV<br>Confusion matrices |  |  |  | 5-Fold CV<br>Confusion matrices |  |  |  | 2-Fold CV<br>Confusion matrices |  |  |  |
| --- | --- | --- | --- | --- | --- | --- | --- | --- | --- | --- | --- | --- |
|  |  | AP | P |  |  | AP | P |  |  | AP | P |  |
| Fold 1 | AP | 87 | 2 | MCC:0.85<br>ACC:0.95 | AP | 175 | 5 | MCC:0.83<br>ACC:0.94 | AP | 436 | 13 | MCC:0.83<br>ACC:0.94 |
|  | P | 4 | 22 |  | P | 8 | 43 |  | P | 13 | 113 |  |
| Fold 2 | AP | 87 | 2 | MCC:0.85<br>ACC:0.95 | AP | 176 | 3 | MCC:0.91<br>ACC:0.97 | AP | 439 | 9 | MCC:0.91<br>ACC:0.97 |
|  | P | 4 | 22 |  | P | 4 | 47 |  | P | 15 | 111 |  |
| Fold 3 | AP | 89 | 1 | MCC:0.92<br>ACC:0.97 | AP | 176 | 4 | MCC:0.91<br>ACC:0.97 |  |  |  |  |
|  | P | 2 | 23 |  | P | 3 | 47 |  |  |  |  |  |
| Fold 4 | AP | 89 | 1 | MCC:0.92<br>ACC:0.97 | AP | 180 | 0 | MCC:0.90<br>ACC:0.97 |  |  |  |  |
|  | P | 2 | 23 |  | P | 8 | 42 |  |  |  |  |  |
| Fold 5 | AP | 86 | 4 | MCC:0.85<br>ACC:0.95 | AP | 175 | 4 | MCC:0.90<br>ACC:0.97 |  |  |  |  |
|  | P | 2 | 23 |  | P | 4 | 46 |  |  |  |  |  |
| Fold 6 | AP | 89 | 1 | MCC:0.95<br>ACC:0.98 |  |  |  |  |  |  |  |  |
|  | P | 1 | 24 |  |  |  |  |  |  |  |  |  |
| Fold 7 | AP | 89 | 1 | MCC:0.81<br>ACC:0.94 |  |  |  |  |  |  |  |  |
|  | P | 6 | 19 |  |  |  |  |  |  |  |  |  |
| Fold 8 | AP | 90 | 0 | MCC:0.95<br>ACC:0.98 |  |  |  |  |  |  |  |  |
|  | P | 2 | 23 |  |  |  |  |  |  |  |  |  |
| Fold 9 | AP | 88 | 2 | MCC:0.84<br>ACC:0.95 |  |  |  |  |  |  |  |  |
|  | P | 4 | 21 |  |  |  |  |  |  |  |  |  |
| Fold 10 | AP | 86 | 3 | MCC:0.90<br>ACC:0.96 |  |  |  |  |  |  |  |  |
|  | P | 1 | 24 |  |  |  |  |  |  |  |  |  |
| Mean values |  |  |  | MCC:0.86<br>ACC:0.96 |  |  |  | MCC:0.89<br>ACC:0.96 |  |  |  | MCC:0.87<br>ACC:0.96 |

**Figure 4:**
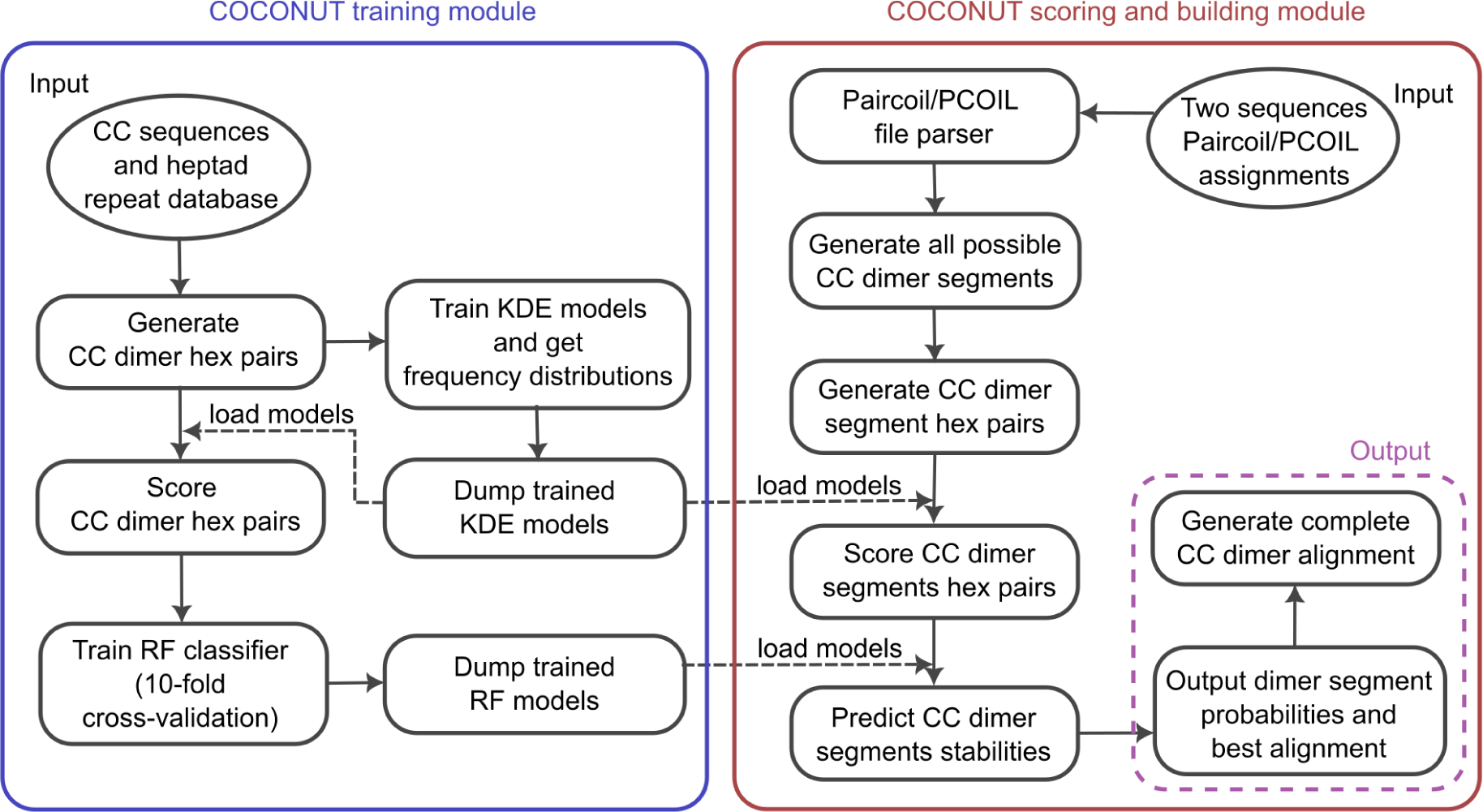
COCONUT algorithm and dataflow. The COCONUT software consists of two modules: training (left) and scoring (right). The training module takes a database of coiled-coil sequences as inputs, along with their heptad repeat assignments. It then generates hex pairs from these sequences, scores them, and uses this information to train and save Kernel Density Estimators and Random Forest models. The scoring module receives two sequences with Paircoil or PCOIL heptad repeat assignments. This data is parsed and generates all potential coiled-coil dimer segments and their corresponding hexpairs. It then scores these segments and predicts their likelihood using the pre-trained models. The final output is a comprehensive list of alignments of the coiled-coil dimers, including probabilities for segment stabilities and optimal alignment between two input sequences.

### 2.5 Building the *n-stranded* coiled-coil models

COCONUT also provides a mathematical framework for constructing *n-stranded* coiled-coil structures. The locus of a coiled-coil consists of two minor helices, each having an axis that winds around a central superhelix (major helix). **Equations 5-7** define the parametric position of a point along the major and minor helices in the coiled-coil structure.

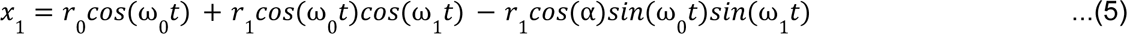

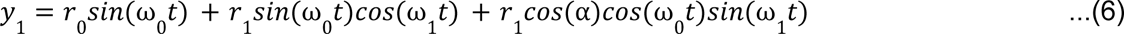

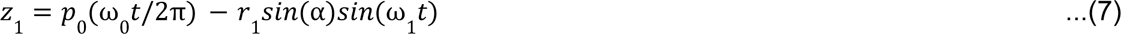

Where *r*_0_ and *r*_1_ is the radius of the major and minor helix, ω_0_ and ω_1_ ω is the angular _1_ frequency of the major and minor helix, α is the phase angle between the major and minor helix, *t* is the parameter that traces the helix (13, 15). For *n-stranded* configurations, the angular frequency of the major helix for each strand is phased by 2π(*strand_id_* − 1)/*n* to account for the varying positions of individual strands, with values of *strand_id_* ranging from 1 to *n*. Additionally, the angular frequency for minor helix in equations 1 and 2 is phased by *heptad_id_* * (4π/7) + 2π/7 to appropriate the position of the C^ɑ^ atom in the coiled-coil with *heptand_id_* = 1 for heptad d, *htptad_id_* = 2 for heptad e, and so on.

Collectively, these equations facilitate the precise generation of a coiled-coil model. The generated models were benchmarked by comparing the models with the atomic coordinates from high-resolution coiled-coil structures from the PDB database (**Fig. 5**).

**Figure 5:**
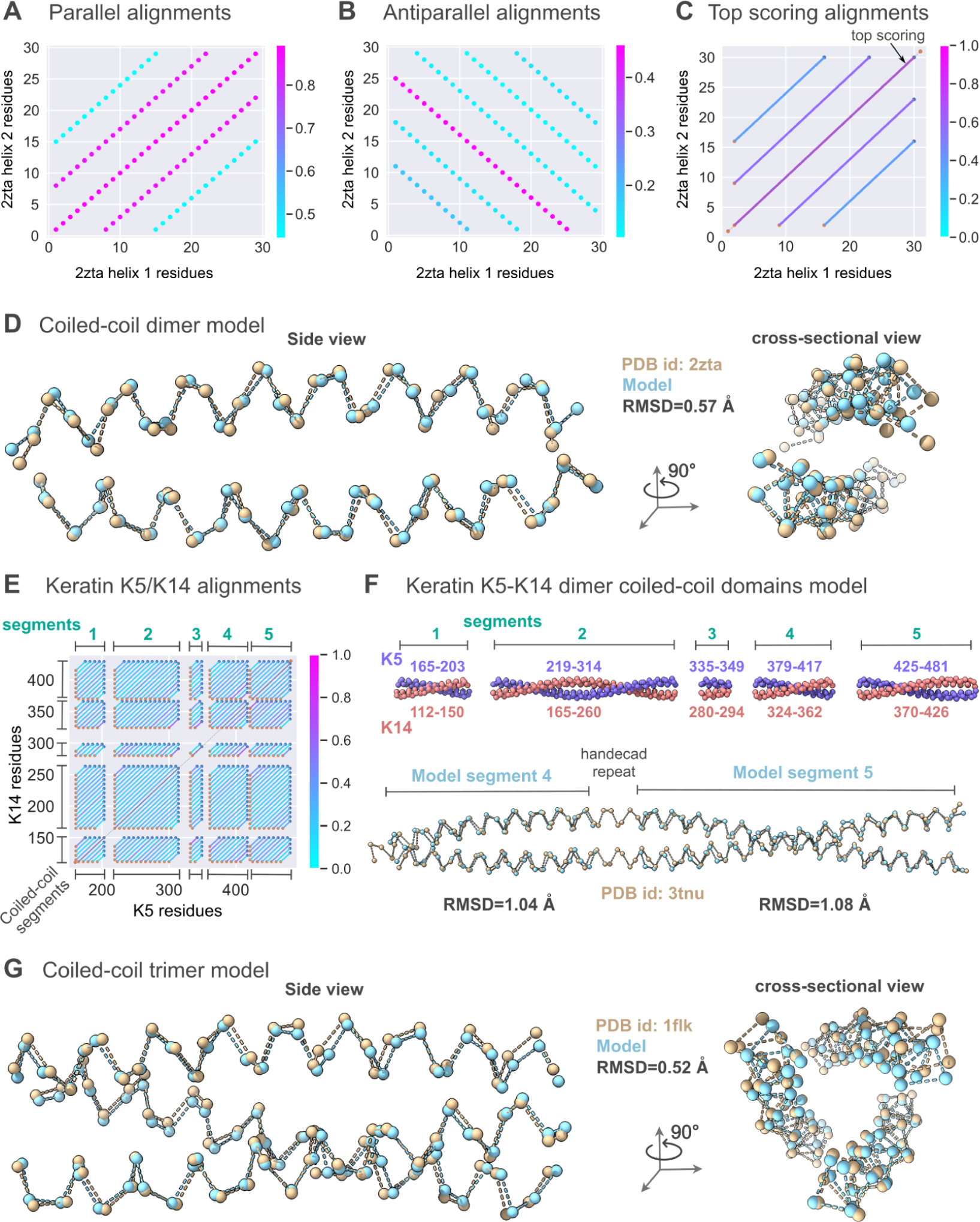
Comparison of COCONUT-generated model and crystal structures. (A, B) All possible parallel and antiparallel alignments of leucine zipper sequences (PDB 2ZTA), with their corresponding alignment scores between 0 and 1 (color bar). (C) Top-scoring alignments for leucine zipper sequences (solid gray line on the diagonal). (D) Side and cross-sectional view of a COCONUT-generated coiled-coil dimer model (cyan) superimposed on the crystal structure (PDB 2ZTA, tan) The RMSD between the model and X-ray structure is 0.57 Å. (E) Possible alignments of the Keratin K5 and K14 sequences coiled-coil domains excluding unstructured head/tail/linker domains. Each alignment suggests a possible association between different K5/K14 coiled-coil segments (turquoise) with color, indicating their interaction strength or compatibility (color bar). Top scoring alignments for K5/K14 sequences with the best alignment (solid gray line) were used for building the K5-K14 dimer model. (F) The K5-K14 dimer model (violet and brick red) with coiled-coil domains numbered from 1-5 (turquoise) indicated by residue numbers in respective colors. The comparison of the model segments 4 and 5 (cyan) with the available crystal structure of resolution 3.00 Å (PDB 3TNU, tan) revealed an RMSD of 1.04 Å and 1.08 Å, respectively. (G) Side and cross-sectional view of a COCONUT-generated coiled-coil trimer model (cyan) superimposed on the crystal structure (PDB 1FLK, tan) revealed an RMSD of 0.52 Å.

## 3 Results

We develop the tool **CO**iled-**CO**il a**N**alysis **UT**ility (COCONUT) to predict the optimal pairing between coiled-coil forming sequences, including their parallel or antiparallel orientation. COCONUT has been trained on a database of 1149 coiled-coil structures that includes 252 parallel and 897 anti-parallel dimers with lengths between 11 to 100 amino acids (**Fig. 1**). The coiled-coil sub-segments consisting of 6 amino acids from each helix (hex pairs) were extracted using a sliding window approach (**Fig. 1C**). A total of 1516 parallel and 3424 anti-parallel hex pairs were identified, providing a comprehensive set for analyzing their molecular interactions.

### 3.1 Hydrophobic contacts serve as the foundation for structural specificity

Hydrophobic core and charge-charge interactions are the two major molecular contacts between helices in a coiled-coil (20). The hydrophobic core is located on the central axis along the length of the coiled-coil and is formed by residues at heptad positions “a” and “d” (**Fig. 1A**). The charge-charge interactions are between residues at heptad positions “e” and “g” that surround the hydrophobic core. Together these molecular contacts form knob-into-hole packing (12, 16) (**Fig. 1B**). Residues at heptad positions “b”, “c”, and “f” do not interact with the neighboring helices and are thus excluded from the analysis. We extracted information about the Knob-into-hole packing residue pairs and found significant differences between parallel and antiparallel dimers.

Specifically, parallel dimers a-a’ heptad pairs can accommodate both branched (at C^β^ positions) and non-branched residues (such as isoleucine, leucine, valine, lysine, and asparagine). These residue pairs account for 41% of all a-a’ pairs. However, d-d’ heptad pairs can accommodate only linear side chain residues (such as leucine) that alone accounts for 36% of all d-d’ pairs (**Fig. 2A**). In antiparallel dimers, a mixed proportion is seen at a-d’ heptad pairs, except for branched side chains amino acids (such as isoleucine) (**Fig. 2B**). The different residue pairings in the hydrophobic core indicates the molecular basis for specificity between helices in the coiled-coils.

The probability distributions (**Fig. 2**) of other heptad pairs (a-g’, d-e’, e-g’) in parallel and (a-e’, d-g’, e-e’, g-g’) in antiparallel dimers also have significantly distinct interaction patterns. However, visual inspection makes these patterns less obvious and needs further analysis to extract information for further use.

### 3.2 Hydrophobic core densities vary between parallel and antiparallel dimer

A stable coiled-coil should maintain a uniform density of atoms in its hydrophobic core (i.e., avoiding volumes of both low or high densities) to avoid exposure to solvents. For a comprehensive interpretation of the hydrophobic core, we consider succeeding and preceding heptad pairs by computing the atomic density scores (ADS) for the “ada”/“dad” hex pairs (**Eq. 1-4**); the ADS scores consider “a”/“d” pairwise scores along with succeeding and preceding “a”/“d” pairwise scores. The ADS scores compute the local deviations in the number and types of residues, which are important for the Knob-into-hole packing (**Fig. 1A, B**). A kernel density estimator was used to model the ADS distributions for both parallel and anti-parallel dimers. The kernel density estimator models were trained to ensure accuracy without over-smoothing, using ‘scott’ method for bandwidth selection (56). The mean of the “ada” distribution is below zero for parallel dimers, whereas for the “dad” distribution, the mean is above zero. This suggests that a low-density region at d-d positions is compensated by a high-density region at a-a positions, making the dimer locally stable by having specific interacting residue pairs. For antiparallel dimers, both “ada” and “dad” hex pairs show identical distributions with a mean around zero owing to their identical a-d/d-a pairings, suggesting uniform density along the length of the dimer. The mean values of the ADS probability distributions of the parallel dimers are statistically different (two-tailed T-test with 95% confidence interval) from the antiparallel dimers (**Fig. 1F**), suggesting the presence of critical information about the distinct structural interaction patterns in coiled-coil dimers. The comparison of ADS probability distributions for parallel and antiparallel dimers suggests that uniform density is maintained across the length of the dimer. In parallel dimers, this occurs as a series of low-high-low densities, whereas in antiparallel dimers, it is a constant density.

### 3.3 COCONUT’s machine learning models and validations

A machine learning approach with Random Forest was utilized to use this information to predict the structural integrity of the coiled-coil forming sequences. A stringent training and validation process was undertaken using stratified 2/5/10-fold cross-validations. Their Matthew’s Correlation coefficients (MCC) are comparable with a value of 0.87, 0.89, and 0.88 for 2/5/10-fold cross-validation, respectively (**Fig. 3A-C**). The consistent results from different cross-validations of the trained models suggest their robustness with unseen data. These models were subsequently prepared for deployment in stability prediction tasks. The models were also evaluated for prediction accuracy across different dimer lengths. The accuracy remains consistent and does not show any trend with dimer length, except for fluctuations due to the decreasing sample size (**Fig. 3D**). We also tested our Random Forest models using the data from dimers that were not part of the training dataset. For antiparallel dimers, 880 out of 897 were predicted correctly, whereas, for parallel dimers, 224 out of 252 were predicted correctly (**Fig. 3E**). A lot of the incorrect predictions are very close to the cutoff probability value of 0.5, suggesting minor tweaks in the methods might improve the performance (**Fig. 3E**). These results suggest the Random Forest models are effective in predicting coiled-coil stability and thus can be utilized for several tasks related to coiled-coils.

### 3.4 COCONUT’s pipeline for coiled-coil analysis and modeling

We developed an automated pipeline leveraging COCONUT predictive scores to analyze the compatibility of coiled-coil forming sequences and construct structural models (**Fig. 4**). The pipeline consists of two modules. The first module is the core of COCONUT, where models are trained and stored for subsequent use (**Fig. 4, left**). The second is the scoring and building module, which utilizes the trained models to evaluate sequences and build structural models (**Fig. 4, right**). Initially, the training module incorporates a coiled-coil sequences database and their heptad repeat annotations. From this database, the software generates coiled-coil dimer hex pairs that serve as the basis for the scoring system. This information guides the training of Kernel Density Estimators and Random Forest classifiers, whose models are retained for future applications. The scoring module takes a pair of sequences with their respective Paircoil/PCOIL assignments and processes this data to identify all possible hex pairs. These segments are then evaluated using the previously trained Kernel Density Estimators models, followed by assessing the stability of each segment using the Random Forest classifier. The pipeline generates a detailed alignment output that proposes the most stable coiled-coil dimer model and assigns probability scores for individual segment’s stability. The output alignment, in turn, facilitates the construction of structural models.

By leveraging this pipeline, we demonstrated the capability of COCONUT through three distinct coiled-coil modeling examples (**Fig. 5**); 1) Identifying the orientation of coiled-coil helices, 2) Identifying the correct coiled-coil helices partners out of many possibilities, and 3) constructing 3D structure of a *n-stranded* coiled-coil structure.

#### 3.4.1 Predicting the coiled-coil orientation in a substitution-sensitive coiled-coil dimer

First, we tested COCONUT’s ability to identify the orientation of the coiled-coil helices using a leucine zipper (PDB 2ZTA), a standard canonical parallel coiled-coil dimer. Previous studies show that GCN4 leucine zipper forms an antiparallel trimer and other parallel higher-order oligomers due to various amino acid substitutions (41, 42, 61). Without assuming any previous knowledge, we used the GCN4 leucine zipper sequence to generate parallel (**Fig. 5A**) and antiparallel (**Fig. 5B**) alignments. The COCONUT scores correctly predicted a parallel dimer with a probability of 0.9 and an antiparallel dimer with a probability of 0.1. The top-scoring parallel alignment (**Fig. 5C**) was used to construct a structural model that resembles the crystal structure with an RMSD of 0.57 Å over 62 heavy atoms (**Fig. 5D**).

#### 3.4.2 Identifying the residue pairings in non-canonical coiled-coil heterodimers

In the second example, we chose a non-canonical coiled-coil that was not part of the training dataset. We used Keratin K5 and K14 intermediate filament sequences as they have been shown to form coiled-coil heterodimers. Based on the sequence analysis, the Keratin K5-K14 heterodimer has long coiled-coil segments connected by linker residues. There are no experimentally derived structures except for small coiled-coil segments. We start with the Keratin K5 (Uniprot ID: P13647) and K14 (Uniprot ID: P02533) sequences to generate all possible coiled-coil segment alignments with COCONUT scores (**Fig. 5E**). Following, the top-scoring alignment is used to build structural models of the coiled-coil regions (**Fig. 5E, F**). We obtained five coiled-coil segments (**Fig. F**). The residues K5:421-424 and K14:366-369 follow a noncanonical heptad repeat known as hendecad repeat. Thus, this region was not modeled, and the resulting coiled-coil was segregated into two parts. We compared the models of these two coiled-coil segments against the available crystal structures (PDB: 3TNU). We find that COCONUT alignment correctly identified the residue pairing between coiled-coil helices, and the corresponding structural model exhibited a strong correspondence with their respective crystal structures with RMSD of 1.04 Å and 1.08 Å, over 63 and 104 heavy atoms.

#### 3.4.3 Building *n-stranded* coiled-coil dimer model

The third example highlights the COCONUT’s proficiency in modeling *n-stranded* coiled-coil structures. In this example, we do not identify the alignment between coiled-coil residues and provide coiled-coil parameters as an input to the pipeline for constructing the structural models. COCONUT provides ready-to-use coiled-coil parameter input files for different types of coiled-coils. For the *n-stranded* coiled-coil, we choose a coiled-coil trimer domain (n=3) of Tumor necrosis factor receptors (PDB 1flk). We extracted its sequence from its structure file and built its model. The model closely matched the crystal structure with an RMSD of 0.52 Å, over 96 heavy atoms.

## 4 Discussion

We present the COCONUT tool, which uses coiled-coil motifs extracted from the CC+ database, predicts the stability and orientation of coiled-coil dimers, and provides a framework for constructing *n-stranded* coiled-coil structural models. The coiled-coil motifs used to build our machine-learning models exhibit varying lengths and maintain a sequence identity below 35%, forming a diverse dataset. COCONUT’s analysis initiates by identifying “hex pairs” within these motifs (**Fig. 1C**), which are crucial for coiled-coil stability. Subsequently, the study examines diverse structural aspects and interactions, including amino acid preferences at heptad positions (**Fig. 1F, 2**).

The role of hydrophobic contacts emerges as a central theme in determining the specificity of coiled-coil structures. Our analysis distinguished between parallel and antiparallel dimers based on differences in residue pair distributions within the hydrophobic core (**Fig. 2**). This study also identified critical variations of hydrophobic contacts essential for maintaining structural integrity. Calculation of atomic density scores from hex pairs allowed insight into density fluctuations within the core, emphasizing the significance of uniformity to prevent destabilization (**Fig. 1F**).

The Random Forest classifier within COCONUT accurately predicted the stability and orientation of coiled-coil dimers (**Fig. 3A-C**). Cross-validation methods showed consistent predictive performance, indicating the tool’s reliability above 96%. Its efficacy was further validated by testing on systems absent from the training dataset, showcasing COCONUT’s proficiency in differentiating parallel and antiparallel dimers (**Fig. 3D**). The utility of COCONUT was showcased through its ability to not only analyze coiled-coil sequences but also to construct accurate structural models. Through various applications, including the prediction of dimer orientation, COCONUT emerged as a robust computational framework for interpreting and modeling coiled-coil structures (**Fig. 5**).

COCONUT is distinguished by its quantitative approach to analyzing coiled-coil interactions, which are essential for understanding protein-protein interactions, as well as designing competitive inhibitors to coiled-coil proteins. COCONUT’s unique feature is the incorporation of machine learning, particularly a Random Forest classifier, to analyze molecular determinants, which enables a prediction of coiled-coil stability and dimer orientation.

Several web servers and software exist for predicting coiled-coil motifs. However, the prediction of complementary coiled-coil motif sequences was limited by the experimental evidence. COCONUT’s approach overcomes these limitations by not only predicting whether specific sequences will form coiled-coils but also by determining their predicted stability and orientation. This information is crucial for accurately modeling protein structures and interpreting protein function within the cellular environment. Moreover, COCONUT could contribute to the understanding of how mutations within these domains could affect overall protein stability, paving the way for insights into disease mechanisms and potential therapeutic targets.

COCONUT’s output has been validated against known structures (**Fig. 5**), demonstrating its ability to construct accurate models of complex protein assemblies, which would have been significantly more challenging without this tool. COCONUT streamlines the traditional modeling approach, eliminating the need for time-consuming trial-and-error methods. One such example is constructing several coiled-coil dimers for a pair of sequences and then identifying the most stable coiled-coil after performing MM-PBSA calculations.

Despite its current capabilities, COCONUT’s application is restricted to dimeric coiled-coils. Expanding its database to encompass higher-order oligomers could reveal more complex interaction patterns broadening its utility. Incorporating evolutionary and contextual data in the COCONUT could enhance its predictive modeling capabilities and refine the accuracy and scope of the predictions.

Future research should focus on enriching the COCONUT database with more varied coiled-coil structures, including those involved in pathological conditions such as intermediate filaments. The role of single missense mutations in coiled coils leading to severe pathological conditions is well documented and provides a fertile ground for therapeutic interventions. Additionally, exploring the interplay between coiled-coil motifs and other protein domains may uncover new functional insights, relevant for the expanding field of synthetic biology.

## Conflict of interest

None declared

## Data availability

The source code of COCONUT is available for free download from https://github.com/neeleshsoni21/COCONUT. The documentation is available in the source code and here: https://neeleshsoni21.github.io/COCONUT/

## Acknowledgments

We would like to thank A. Sali, for insightful discussions, and Sali lab members at UCSF; I. Echeverria, A. Zalevsky, A. Latham, and K. Huang, for their extensive comments on the manuscript. We would also like to thank members of the COSPI lab at IISER Pune for valuable discussions.

## Notes

### Competing Interest Statement

The authors have declared no competing interest.

## References

1. Hartmann, M.D. (2017) Functional and Structural Roles of Coiled Coils. In Parry, D.A.D., Squire, J.M. (eds), Fibrous Proteins: Structures and Mechanisms. Springer International Publishing, Cham, pp. 63–93.

2. Rackham, O.J.L., Madera, M., Armstrong, C.T., Vincent, T.L., Woolfson, D.N. and Gough, J. (2010) The evolution and structure prediction of coiled coils across all genomes. J. Mol. Biol., 403, 480–493.

3. Lupas, A. (1996) Coiled coils: new structures and new functions. Trends Biochem. Sci., 21, 375–382.

4. Burkhard, P., Stetefeld, J. and Strelkov, S.V. (2001) Coiled coils: a highly versatile protein folding motif. Trends Cell Biol., 11, 82–88.

5. Weichsel, A., Kievenaar, J.A., Curry, R., Croft, J.T. and Montfort, W.R. (2019) Instability in a coiled-coil signaling helix is conserved for signal transduction in soluble guanylyl cyclase. Protein Sci., 28, 1830–1839.

6. Schweke, H., Pacesa, M., Levin, T., Goverde, C.A., Kumar, P., Duhoo, Y., Dornfeld, L.J., Dubreuil, B., Georgeon, S., Ovchinnikov, S., et al. (2024) An atlas of protein homo-oligomerization across domains of life. Cell, 187, 999–1010.e15.

7. Parry, D.A.D. (2014) Fifty years of fibrous protein research: a personal retrospective. J. Struct. Biol., 186, 320–334.

8. Truebestein, L. and Leonard, T.A. (2016) Coiled-coils: The long and short of it. Bioessays, 38, 903–916.

9. Park, W.M. (2020) Coiled-Coils: the Molecular Zippers that Self-Assemble Protein Nanostructures. Int. J. Mol. Sci., 21.

10. Lapenta, F., Aupič, J., Vezzoli, M., Strmšek, Ž., Da Vela, S., Svergun, D.I., Carazo, J.M., Melero, R. and Jerala, R. (2021) Self-assembly and regulation of protein cages from pre-organised coiled-coil modules. Nat. Commun., 12, 939.

11. Lupas, A.N. and Bassler, J. (2017) Coiled Coils - A Model System for the 21st Century. Trends Biochem. Sci., 42, 130–140.

12. Crick, F.H.C. (1952) Is alpha-keratin a coiled coil? Nature, 170, 882–883.

13. Crick, F.H.C. (1953) The Fourier transform of a coiled-coil. Acta Crystallogr., 6, 685–689.

14. Parry, D.A.D., Fraser, R.D.B. and Squire, J.M. (2008) Fifty years of coiled-coils and alpha-helical bundles: a close relationship between sequence and structure. J. Struct. Biol., 163, 258–269.

15. Crick, F.H.C. and IUCr (1953) The packing of -helices: simple coiled-coils. Acta Crystallogr., 6, 689–697.

16. Woolfson, D.N. (2023) Understanding a protein fold: The physics, chemistry, and biology of α-helical coiled coils. J. Biol. Chem., 299, 104579.

17. Strauss, H.M. and Keller, S. (2008) Pharmacological Interference with Protein-Protein Interactions Mediated by Coiled-Coil Motifs. In Klussmann, E., Scott, J. (eds), Protein-Protein Interactions as New Drug Targets. Springer Berlin Heidelberg, Berlin, Heidelberg, pp. 461–482.

18. Szczepaniak, K., Bukala, A., da Silva Neto, A.M., Ludwiczak, J. and Dunin-Horkawicz, S. (2021) A library of coiled-coil domains: from regular bundles to peculiar twists. Bioinformatics, 36, 5368–5376.

19. McDonnell, A.V., Jiang, T., Keating, A.E. and Berger, B. (2006) Paircoil2: improved prediction of coiled coils from sequence. Bioinformatics, 22, 356–358.

20. Ludwiczak, J., Winski, A., Szczepaniak, K., Alva, V. and Dunin-Horkawicz, S. (2019) DeepCoil-a fast and accurate prediction of coiled-coil domains in protein sequences. Bioinformatics, 35, 2790–2795.

21. Hewagama, N.D., Uchida, M., Wang, Y., Kraj, P., Lee, B. and Douglas, T. (2023) Higher-order VLP-based protein macromolecular framework structures assembled via coiled-coil interactions. Biomacromolecules, 10.1021/acs.biomac.3c00410.

22. Brinkmann, D., Nandoor, S., Kalita, J., Tripet, B. and Hodges, R. (2004) CoCoLysis : A web-accessible coiled-coil protein database with analysis tools. on Bioinformatics and *…*.

23. Kumar, P., Petrenas, R., Dawson, W.M., Schweke, H., Levy, E.D. and Woolfson, D.N. (2023) CC+ : A searchable database of validated coiled coils in PDB structures and AlphaFold2 models. Protein Sci., 32, e4789.

24. Lupas, A., Van Dyke, M. and Stock, J. (1991) Predicting coiled coils from protein sequences. Science, 252, 1162–1164.

25. Vincent, T.L., Green, P.J. and Woolfson, D.N. (2013) LOGICOIL--multi-state prediction of coiled-coil oligomeric state. Bioinformatics, 29, 69–76.

26. Lupas, A. (1996) Prediction and analysis of coiled-coil structures. Methods Enzymol., 266, 513–525.

27. Gruber, M., Söding, J. and Lupas, A.N. (2006) Comparative analysis of coiled-coil prediction methods. J. Struct. Biol., 155, 140–145.

28. Guzenko, D. and Strelkov, S.V. (2018) CCFold: rapid and accurate prediction of coiled-coil structures and application to modelling intermediate filaments. Bioinformatics, 34, 215–222.

29. Madeo, G., Savojardo, C., Manfredi, M., Martelli, P.L. and Casadio, R. (2023) CoCoNat: a novel method based on deep learning for coiled-coil prediction. Bioinformatics, 39.

30. Feng, S.-H., Xia, C.-Q. and Shen, H.-B. (2022) CoCoPRED: coiled-coil protein structural feature prediction from amino acid sequence using deep neural networks. Bioinformatics, 38, 720–729.

31. Walshaw, J. and Woolfson, D.N. (2001) Socket: a program for identifying and analysing coiled-coil motifs within protein structures. J. Mol. Biol., 307, 1427–1450.

32. Delorenzi, M. and Speed, T. (2002) An HMM model for coiled-coil domains and a comparison with P\SSM-based predictions. Bioinformatics, 18, 617–625.

33. Simm, D., Hatje, K., Waack, S. and Kollmar, M. (2021) Critical assessment of coiled-coil predictions based on protein structure data. Sci. Rep., 11, 12439.

34. Apostolovic, B., Danial, M. and Klok, H.-A. (2010) Coiled coils: attractive protein folding motifs for the fabrication of self-assembled, responsive and bioactive materials. Chem. Soc. Rev., 39, 3541–3575.

35. Fariselli, P., Molinini, D., Casadio, R. and Krogh, A. (2007) Prediction of Structurally-Determined Coiled-Coil Domains with Hidden Markov Models. In Bioinformatics Research and Development. Springer Berlin Heidelberg, pp. 292–302.

36. Parry, D.A. (1982) Coiled-coils in alpha-helix-containing proteins: analysis of the residue types within the heptad repeat and the use of these data in the prediction of coiled-coils in other proteins. Biosci. Rep., 2, 1017–1024.

37. Karami, Y., Saighi, P., Vanderhaegen, R., Gerlier, D., Longhi, S., Laine, E. and Carbone, A. (2020) Predicting substitutions to modulate disorder and stability in coiled-coils. BMC Bioinformatics, 21, 573.

38. Mason, J.M. and Arndt, K.M. (2004) Coiled coil domains: stability, specificity, and biological implications. Chembiochem, 5, 170–176.

39. Yu, Y.B. (2002) Coiled-coils: stability, specificity, and drug delivery potential. Adv. Drug Deliv. Rev., 54, 1113–1129.

40. Grigoryan, G. and Keating, A.E. (2008) Structural specificity in coiled-coil interactions. Curr. Opin. Struct. Biol., 18, 477–483.

41. Harbury, P.B., Zhang, T., Kim, P.S. and Alber, T. (1993) A switch between two-, three-, and four-stranded coiled coils in GCN4 leucine zipper mutants. Science, 262, 1401–1407.

42. Oshaben, K.M., Salari, R., McCaslin, D.R., Chong, L.T. and Horne, W.S. (2012) The native GCN4 leucine-zipper domain does not uniquely specify a dimeric oligomerization state. Biochemistry, 51, 9581–9591.

43. Lowey, S. (1965) COMPARATIVE STUDY OF THE ALPHA-HELICAL MUSCLE PROTEINS. TYROSYL TITRATION AND EFFECT OF PH ON CONFORMATION. J. Biol. Chem., 240, 2421–2427.

44. Hodges, R.S., Sodek, J., Smillie, L.B. and Jurasek, L. (1973) Tropomyosin: Amino Acid Sequence and Coiled-Coil Structure. Cold Spring Harb. Symp. Quant. Biol., 37, 299–310.

45. Dutta, K., Alexandrov, A., Huang, H. and Pascal, S.M. (2001) pH-induced folding of an apoptotic coiled coil. Protein Sci., 10, 2531–2540.

46. Dutta, K., Engler, F.A., Cotton, L., Alexandrov, A., Bedi, G.S., Colquhoun, J. and Pascal, S.M. (2003) Stabilization of a pH-sensitive apoptosis-linked coiled coil through single point mutations. Protein Sci., 12, 257–265.

47. Liu, Y., Zhou, X., Liu, W. and Miao, W. (2020) The stability of the coiled-coil structure near to N-terminus influence the heat resistance of harpin proteins from Xanthomonas. BMC Microbiol., 20, 344.

48. Mason, J.M., Hagemann, U.B. and Arndt, K.M. (2009) Role of hydrophobic and electrostatic interactions in coiled coil stability and specificity. Biochemistry, 48, 10380–10388.

49. Kirwan, J.P., Kwok, S.C., McReynolds, S., Mills, J., Osguthorpe, D. and Hodges, R.S. (2009) Unique role of clusters of electrostatic attractions in controlling the stability of two-stranded alpha-helical coiled-coils. Adv. Exp. Med. Biol., 611, 77–78.

50. Zhou, N.E., Kay, C.M. and Hodges, R.S. (1994) The net energetic contribution of interhelical electrostatic attractions to coiled-coil stability. Protein Eng., 7, 1365–1372.

51. Parmar, A.S., Joshi, M., Nosker, P.L., Hasan, N.F. and Nanda, V. (2013) Control of Collagen Stability and Heterotrimer Specificity through Repulsive Electrostatic Interactions. Biomolecules, 3, 986–996.

52. Monera, O.D., Zhou, N.E., Lavigne, P., Kay, C.M. and Hodges, R.S. (1996) Formation of Parallel and Antiparallel Coiled-coils Controlled by the Relative Positions of Alanine Residues in the Hydrophobic Core (∗). J. Biol. Chem., 271, 3995–4001.

53. Sen, N., Kanitkar, T.R., Roy, A.A., Soni, N., Amritkar, K., Supekar, S., Nair, S., Singh, G. and Madhusudhan, M.S. (2019) Predicting and designing therapeutics against the Nipah virus. PLoS Negl. Trop. Dis., 13, e0007419.

54. Testa, O.D., Moutevelis, E. and Woolfson, D.N. (2009) CC+: a relational database of coiled-coil structures. Nucleic Acids Res., 37, D315–22.

55. Silverman, B.W. (1986) Density Estimation for Statistics and Data Analysis Estimation Density Kluwer Academic Publishers.

56. Scott, D.W. (1992) Multivariate Density Estimation: Theory, Practice, and Visualization John Wiley & Sons.

57. Jones, E., Oliphant, T., Peterson, P. and Others (2001) SciPy.org. SciPy: Open source scientific tools for Python2.

58. Breiman, L. (2001) Random Forests. Mach. Learn., 45, 5–32.

59. Pedregosa, F., Varoquaux, G., Gramfort, A., Michel, V., Thirion, B., Grisel, O., Blondel, M., Louppe, G., Prettenhofer, P., Weiss, R., et al. (2011) Scikit-learn: Machine Learning in Python. J. Mach. Learn. Res., **abs/**1201.0490.

60. Baldi, P., Brunak, S., Chauvin, Y., Andersen, C.A. and Nielsen, H. (2000) Assessing the accuracy of prediction algorithms for classification: an overview. Bioinformatics, 16, 412–424.

61. Holton, J. and Alber, T. (2004) Automated protein crystal structure determination using ELVES. Proc. Natl. Acad. Sci. U. S. A., 101, 1537–1542.

62. Bjelić, S., Wieser, M., Frey, D., Stirnimann, C.U., Chance, M.R., Jaussi, R., Steinmetz, M.O. and Kammerer, R.A. (2013) Structural basis for the oligomerization-state switch from a dimer to a trimer of an engineered cortexillin-1 coiled-coil variant. PLoS One, 8, e63370.

63. Ciani, B., Bjelic, S., Honnappa, S., Jawhari, H., Jaussi, R., Payapilly, A., Jowitt, T., Steinmetz, M.O. and Kammerer, R.A. (2010) Molecular basis of coiled-coil oligomerization-state specificity. Proc. Natl. Acad. Sci. U. S. A., 107, 19850–19855.

